# Comparative pharmacology of direct oral anticoagulants and vitamin K antagonist

**DOI:** 10.1101/2022.11.10.515974

**Authors:** Arun HS Kumar

**Author notes:** **Correspondence**, Arun HS Kumar, DVM, PhD., Room 216, School of Veterinary Medicine, University College Dublin, Belfield, Dublin-04, Ireland.

## Abstract

**Background:** The prevalence of thrombus and use of anticoagulants is routine in clinical cardiology practice. Vitamin K antagonists (VKA) and/or Direct oral anticoagulants (DOAC) are used for resolution of the thrombus. Despite similar anticoagulation efficacy, use of DOAC is preferred due to their superior safety margin and reduced risk of bleeding. Currently the following DOAC are available for the prevention of thrombosis, i.e., dabigatran, rivaroxaban, apixaban, edoxaban, and betrixaban. This study evaluates the comparative pharmacology of these DOAC and VKA to assess clinical preference.

**Materials and Methods:** The human specific targets of DOAC and VKA (Warfarin) were identified from the SwissTargetPrediction server and analysed for their affinity. The targets were subclassified into functional categories and the relative proportion of each of the functional categories among the total number of targets was estimated. A novel concentration affinity (CA) ratio system was developed for the drugs to assess their safety margin and compared.

**Results:** The following targets were identified has high affinity targets of DOAC or VKA i.e., coagulation factor X, hERG, matriptase, multidrug and toxin extrusion protein 1, plasminogen, quinone reductase 1 & 2, serine protease hepsin, solute carrier family 22 member 2 and thrombin. Apixaban and rivaroxaban were observed to have superior anticoagulation pharmacology compared to the other DOAC or VKA. Edoxaban and betrixaban were observed to have affinity against hERG, which carries the risk of prolonging QT interval and triggering ventricular tachyarrhythmia.

**Conclusion:** This study shows the comparative pharmacology of DOAC and VKA and suggests preferential use of apixaban or rivaroxaban due to their superior pharmacodynamic effects and wider safety margin.

## Introduction

The prevalence of thrombus especially in left ventricle significantly increases the risk of acute ischemic events.^[1–3]^ This risk is further enhanced in presence of comorbidities such as diabetes, hypertension, recent incidence of myocardial infarction (MI) and several lifestyle factors negatively impacting health.^[4–6]^ The incidence of left ventricle thrombus (LVT) is reported to range from 15 to 60% in patients post recovery from MI, while the incidence of LVT in general population without any comorbidities is not known, but is assumed to be low (<5%).^[1, 2, 5, 7]^ The current approaches to managing incidence of LVT include use of vitamin K antagonists (VKA) and/or Direct oral anticoagulants (DOAC) until resolution of the thrombus (~ 3 to 6 months).^[1, 5, 7-9]^ Although recent clinical approaches show preferential use of DOAC over VKA, due to better patient compliance and drug safety issues (reduced bleeding and limited adverse drug interactions).^[10–13]^ Currently following five DOAC are available to be used for the prevention of thrombosis, i.e., dabigatran, rivaroxaban, apixaban, edoxaban, and betrixaban. Several studies have compared the efficacy and safety of different anticoagulant approaches (i.e., DOAC, VKA and Heparin) to achieve resolution of thrombus under different clinical settings,^[12–14]^ however such comparative efficacy of all the available DOAC with reference to their pharmacodynamic effects is lacking. Hence in this study comparative pharmacology of all five DOAC and VKA (warfarin) was assessed.

## Materials and Methods

The targets of DOAC (dabigatran, rivaroxaban, apixaban, edoxaban, and betrixaban) and VKA (Warfarin) were identified from the SwissTargetPrediction server as reported before ^[15, 16]^ and analysed. Briefly, the isomeric SMILES sequence of the drugs obtained from the PubChem database were inputted into the SwissTargetPrediction server to identify the targets specific to homo sapiens. The target list for each of the drugs were processed based on their probability scores to identify highest affinity targets and compared. The targets without any probability score were excluded from the analysis. Drugs showing affinity with multiple targets were further analysed by subclassifying the targets into functional categories and the relative proportion of each of the functional categories among the total number of targets was estimated.

To assess the safety margin of the drugs, the plasma concentration (μM) of the drug achievable following different dose (low, mid and high dose) administration was estimated and the dose dependent concentration affinity ratio (CA ratio) for each of the drugs was calculated. The CA ratio is presented as mean ± SD of the values from low, mid and high dose. The affinity (μM) of the DOAC (to factor Xa/thrombin) and VKA (to Quinone reductase 1) was text mined from the literature. The volume of distribution (L) of the drugs reported in the DrugBank database was used for estimation of the plasma drug concentration (μM) achievable.^[17, 18]^ As the estimated concentration positively correlated with the plasma drug concentration (μM) reported in the literature, it was used for estimation of CA ratio, which reflects the safety margin of drug and the likeliness of significant off target effects when the CA ratio is high.

## Results

The following targets were identified has high affinity targets (probability score >0.8) of DOAC or VKA i.e., coagulation factor X, hERG, matriptase, multidrug and toxin extrusion protein 1, plasminogen, quinone reductase 1 & 2, serine protease hepsin, solute carrier family 22 member 2 and thrombin. Although DOAC showed very high affinity (probability score >0.8) with at least one specific target, in contrast VKA (warfarin) was observed to have weak affinity to all its targets with the maximum affinity (probability score: 0.13) observed for Quinone reductase 1 (Table 1). Among the DOAC, dabigatran was observed to have high affinity (probability score >0.8) for maximum number of targets (coagulation factor X, multidrug/toxin extrusion protein 1, plasminogen, serine protease hepsin, solute carrier family 22 member 2 and thrombin) (Table 1).

**Table 1:** Comparative pharmacology of direct oral anticoagulants and antiplatelet agent against their major targets. Values indicate the probability interaction score of drug with its target.

| TARGETS | Warfarin | Rivaroxaban | Apixaban | Dabigatran | Edoxaban | Betrixaban |
| --- | --- | --- | --- | --- | --- | --- |
| Coagulation Factor X |  | 0.99 | 0.96 | 0.99 | 1 | 1 |
| hERG |  |  |  |  | 0.13 | 1 |
| Matriptase |  | 0.99 |  |  |  |  |
| Multidrug and toxin extrusion protein 1 |  |  |  | 0.99 |  |  |
| Plasminogen |  |  |  | 0.99 |  |  |
| Quinone reductase 1 | 0.13 |  |  |  |  |  |
| Quinone reductase 2 |  |  |  | 0.33 |  |  |
| Serine protease hepsin |  |  |  | 0.99 |  |  |
| Solute carrier family 22 member 2 |  |  |  | 0.99 |  |  |
| Thrombin |  | 0.10 | 0.96 | 0.99 | 0.13 |  |

Based on the analysis profile observed in this study it appears that apixaban has the superior anticoagulation pharmacology compared to the other DOAC or VKA. The superiority of apixaban is due to its specific high affinity to selective targets (coagulation factor X and thrombin) involved in the coagulation cascade and its low affinity (probability score: <0.2) to off targets (tables 1 and 4). Besides apixaban, rivaroxaban also showed high affinity to coagulation factor X and matriptase but it had a lower affinity (probability score: 0.10) to thrombin. However unlike apixaban, rivaroxaban showed highly selective and specific pharmacology against proteases (table 3, figure 1). Edoxaban and betrixaban also showed selective high affinity (probability score: >0.8) towards coagulation factor X, however both these DOAC were observed to have affinity against hERG (table 1), which carries the risk of prolonging QT interval and triggering ventricular tachyarrhythmia.

**Figure 1:**
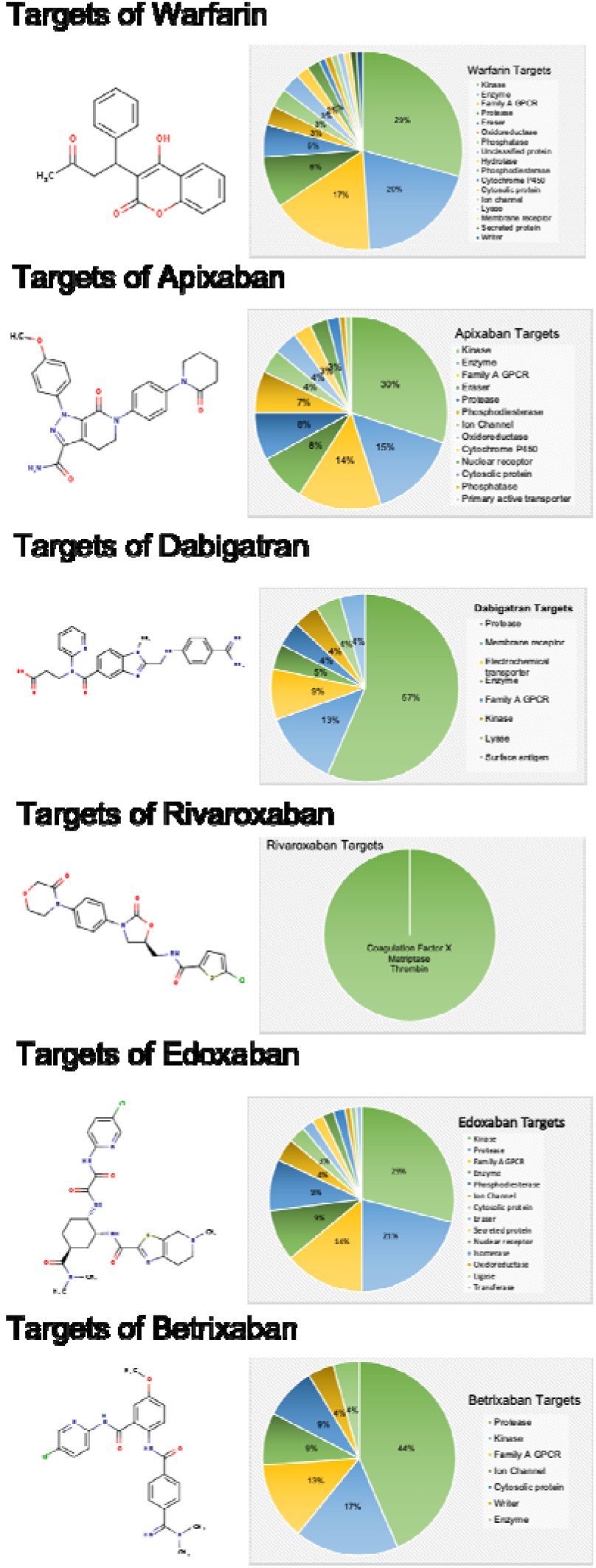
All the potential targets of warfarin, apixaban, dabigatran, rivaroxaban, edoxaban, and betrixaban in humans grouped based on functional categories. The target functional categories showing probability of interaction with the drug are expressed as the percent of the total pool of targets.

**Table 2:**
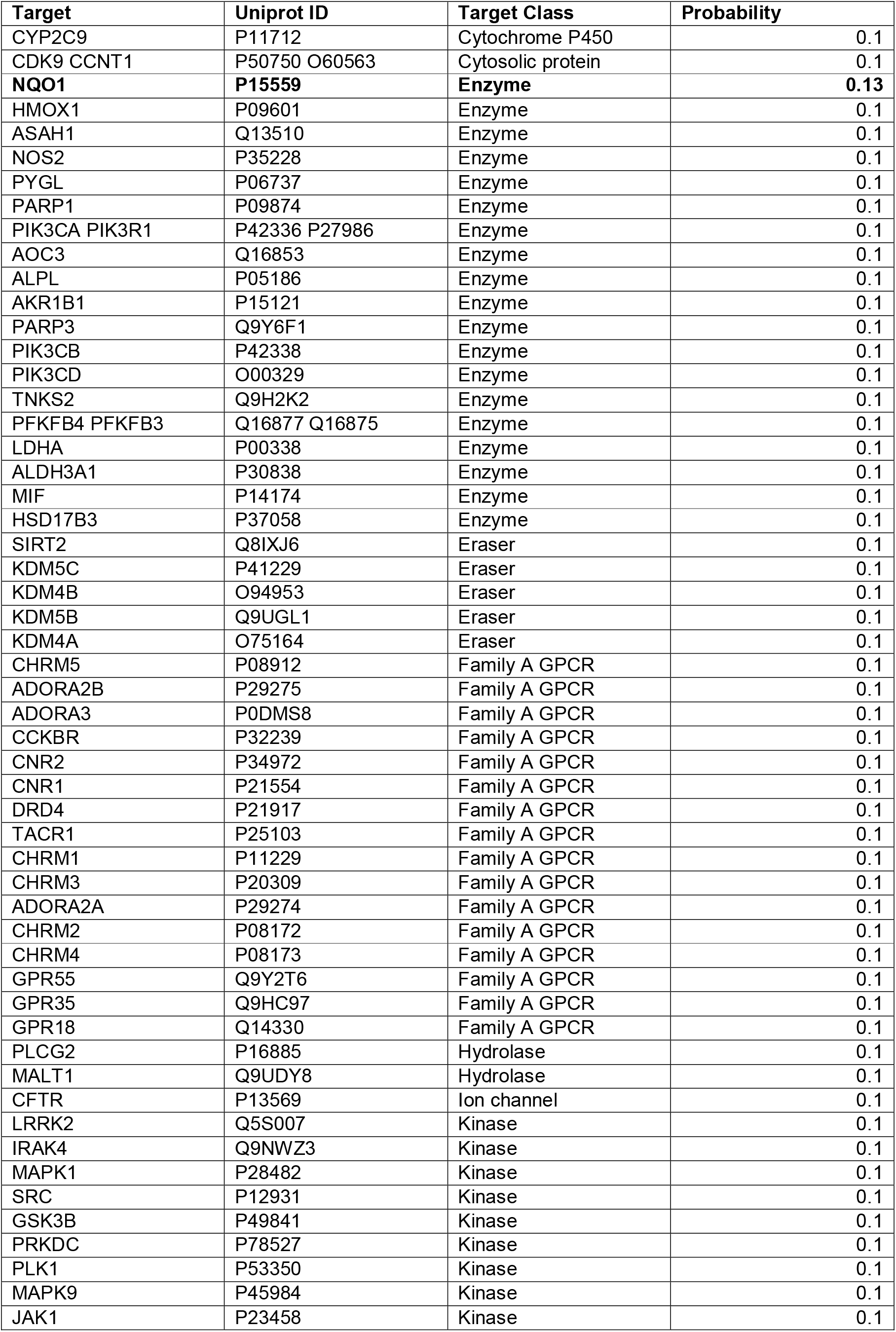
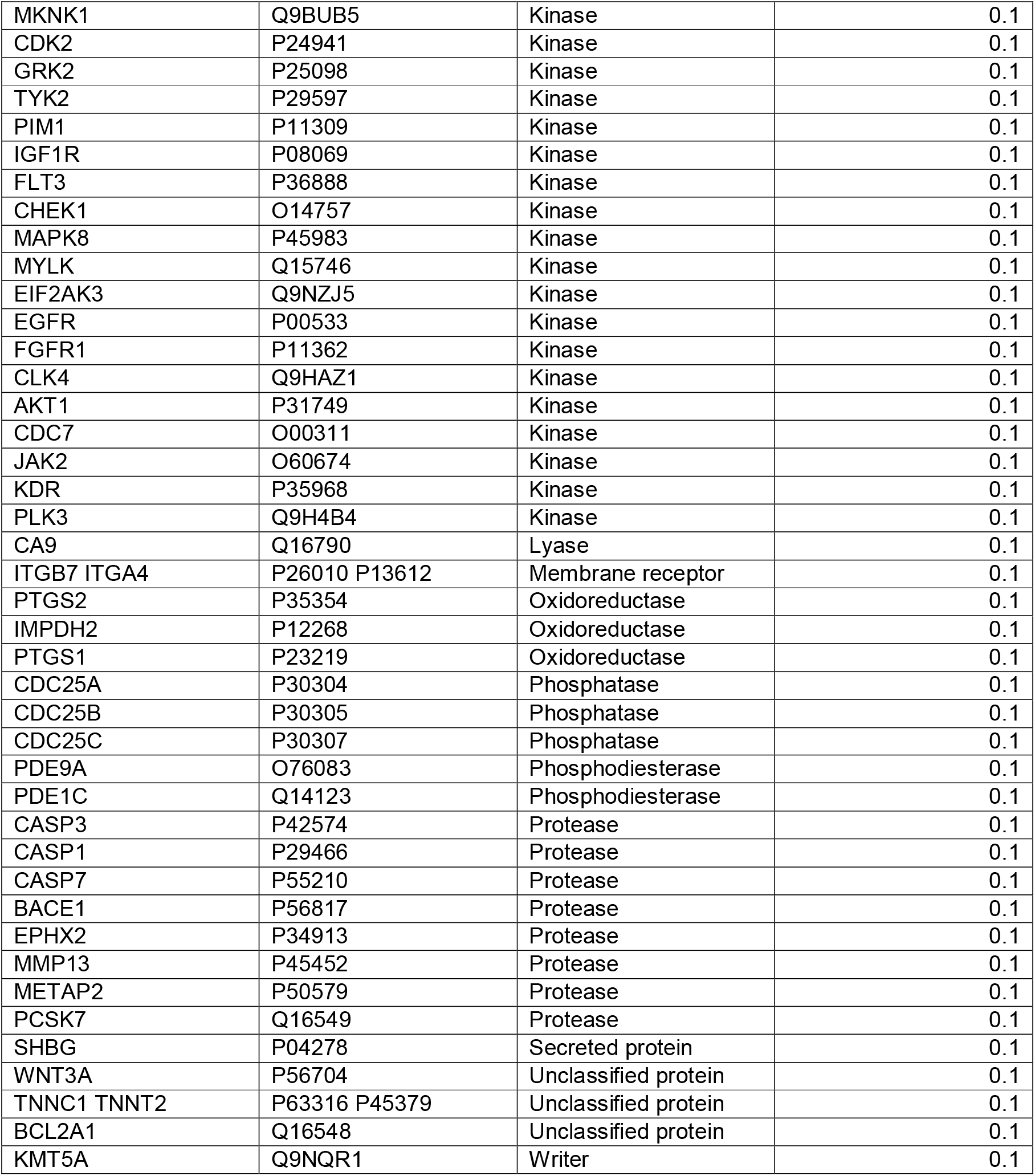
All the potential targets of warfarin in humans. Values indicate the probability interaction score of drug with its target, with value of 1 indicating significant interaction while value of 0.1 indicates weaker interaction.

| Target | Uniprot ID | Target Class | Probability |
| --- | --- | --- | --- |
| CYP2C9 | P11712 | Cytochrome P450 | 0.1 |
| CDK9 CCNT1 | P50750 O60563 | Cytosolic protein | 0.1 |
| **NQO1** | **P15559** | **Enzyme** | **0.13** |
| HMOX1 | P09601 | Enzyme | 0.1 |
| ASAH1 | Q13510 | Enzyme | 0.1 |
| NOS2 | P35228 | Enzyme | 0.1 |
| PYGL | P06737 | Enzyme | 0.1 |
| PARP1 | P09874 | Enzyme | 0.1 |
| PIK3CA PIK3R1 | P42336 P27986 | Enzyme | 0.1 |
| AOC3 | Q16853 | Enzyme | 0.1 |
| ALPL | P05186 | Enzyme | 0.1 |
| AKR1B1 | P15121 | Enzyme | 0.1 |
| PARP3 | Q9Y6F1 | Enzyme | 0.1 |
| PIK3CB | P42338 | Enzyme | 0.1 |
| PIK3CD | O00329 | Enzyme | 0.1 |
| TNKS2 | Q9H2K2 | Enzyme | 0.1 |
| PFKFB4 PFKFB3 | Q16877 Q16875 | Enzyme | 0.1 |
| LDHA | P00338 | Enzyme | 0.1 |
| ALDH3A1 | P30838 | Enzyme | 0.1 |
| MIF | P14174 | Enzyme | 0.1 |
| HSD17B3 | P37058 | Enzyme | 0.1 |
| SIRT2 | Q8IXJ6 | Eraser | 0.1 |
| KDM5C | P41229 | Eraser | 0.1 |
| KDM4B | O94953 | Eraser | 0.1 |
| KDM5B | Q9UGL1 | Eraser | 0.1 |
| KDM4A | O75164 | Eraser | 0.1 |
| CHRM5 | P08912 | Family A GPCR | 0.1 |
| ADORA2B | P29275 | Family A GPCR | 0.1 |
| ADORA3 | P0DMS8 | Family A GPCR | 0.1 |
| CCKBR | P32239 | Family A GPCR | 0.1 |
| CNR2 | P34972 | Family A GPCR | 0.1 |
| CNR1 | P21554 | Family A GPCR | 0.1 |
| DRD4 | P21917 | Family A GPCR | 0.1 |
| TACR1 | P25103 | Family A GPCR | 0.1 |
| CHRM1 | P11229 | Family A GPCR | 0.1 |
| CHRM3 | P20309 | Family A GPCR | 0.1 |
| ADORA2A | P29274 | Family A GPCR | 0.1 |
| CHRM2 | P08172 | Family A GPCR | 0.1 |
| CHRM4 | P08173 | Family A GPCR | 0.1 |
| GPR55 | Q9Y2T6 | Family A GPCR | 0.1 |
| GPR35 | Q9HC97 | Family A GPCR | 0.1 |
| GPR18 | Q14330 | Family A GPCR | 0.1 |
| PLCG2 | P16885 | Hydrolase | 0.1 |
| MALT1 | Q9UDY8 | Hydrolase | 0.1 |
| CFTR | P13569 | Ion channel | 0.1 |
| LRRK2 | Q5S007 | Kinase | 0.1 |
| IRAK4 | Q9NWZ3 | Kinase | 0.1 |
| MAPK1 | P28482 | Kinase | 0.1 |
| SRC | P12931 | Kinase | 0.1 |
| GSK3B | P49841 | Kinase | 0.1 |
| PRKDC | P78527 | Kinase | 0.1 |
| PLK1 | P53350 | Kinase | 0.1 |
| MAPK9 | P45984 | Kinase | 0.1 |
| JAK1 | P23458 | Kinase | 0.1 |
| MKNK1 | Q9BUB5 | Kinase | 0.1 |
| CDK2 | P24941 | Kinase | 0.1 |
| GRK2 | P25098 | Kinase | 0.1 |
| TYK2 | P29597 | Kinase | 0.1 |
| PIM1 | P11309 | Kinase | 0.1 |
| IGF1R | P08069 | Kinase | 0.1 |
| FLT3 | P36888 | Kinase | 0.1 |
| CHEK1 | O14757 | Kinase | 0.1 |
| MAPK8 | P45983 | Kinase | 0.1 |
| MYLK | Q15746 | Kinase | 0.1 |
| EIF2AK3 | Q9NZJ5 | Kinase | 0.1 |
| EGFR | P00533 | Kinase | 0.1 |
| FGFR1 | P11362 | Kinase | 0.1 |
| CLK4 | Q9HAZ1 | Kinase | 0.1 |
| AKT1 | P31749 | Kinase | 0.1 |
| CDC7 | O00311 | Kinase | 0.1 |
| JAK2 | O60674 | Kinase | 0.1 |
| KDR | P35968 | Kinase | 0.1 |
| PLK3 | Q9H4B4 | Kinase | 0.1 |
| CA9 | Q16790 | Lyase | 0.1 |
| ITGB7 ITGA4 | P26010 P13612 | Membrane receptor | 0.1 |
| PTGS2 | P35354 | Oxidoreductase | 0.1 |
| IMPDH2 | P12268 | Oxidoreductase | 0.1 |
| PTGS1 | P23219 | Oxidoreductase | 0.1 |
| CDC25A | P30304 | Phosphatase | 0.1 |
| CDC25B | P30305 | Phosphatase | 0.1 |
| CDC25C | P30307 | Phosphatase | 0.1 |
| PDE9A | O76083 | Phosphodiesterase | 0.1 |
| PDE1C | Q14123 | Phosphodiesterase | 0.1 |
| CASP3 | P42574 | Protease | 0.1 |
| CASP1 | P29466 | Protease | 0.1 |
| CASP7 | P55210 | Protease | 0.1 |
| BACE1 | P56817 | Protease | 0.1 |
| EPHX2 | P34913 | Protease | 0.1 |
| MMP13 | P45452 | Protease | 0.1 |
| METAP2 | P50579 | Protease | 0.1 |
| PCSK7 | Q16549 | Protease | 0.1 |
| SHBG | P04278 | Secreted protein | 0.1 |
| WNT3A | P56704 | Unclassified protein | 0.1 |
| TNNC1 TNNT2 | P63316 P45379 | Unclassified protein | 0.1 |
| BCL2A1 | Q16548 | Unclassified protein | 0.1 |
| KMT5A | Q9NQR1 | Writer | 0.1 |

**Table 3:** All the potential targets of rivaroxaban in humans.

| Target | Uniprot ID | Target Class | Probability* |
| --- | --- | --- | --- |
| **F10** | **P00742** | **Protease** | **0.99** |
| **ST14** | **Q9Y5Y6** | **Protease** | **0.99** |
| F2 | P00734 | Protease | 0.10 |

\*Values indicate the probability interaction score of drug with its target, with value of 1 indicating significant interaction while value of 0.1 indicates weaker interaction.

**Table 4:**
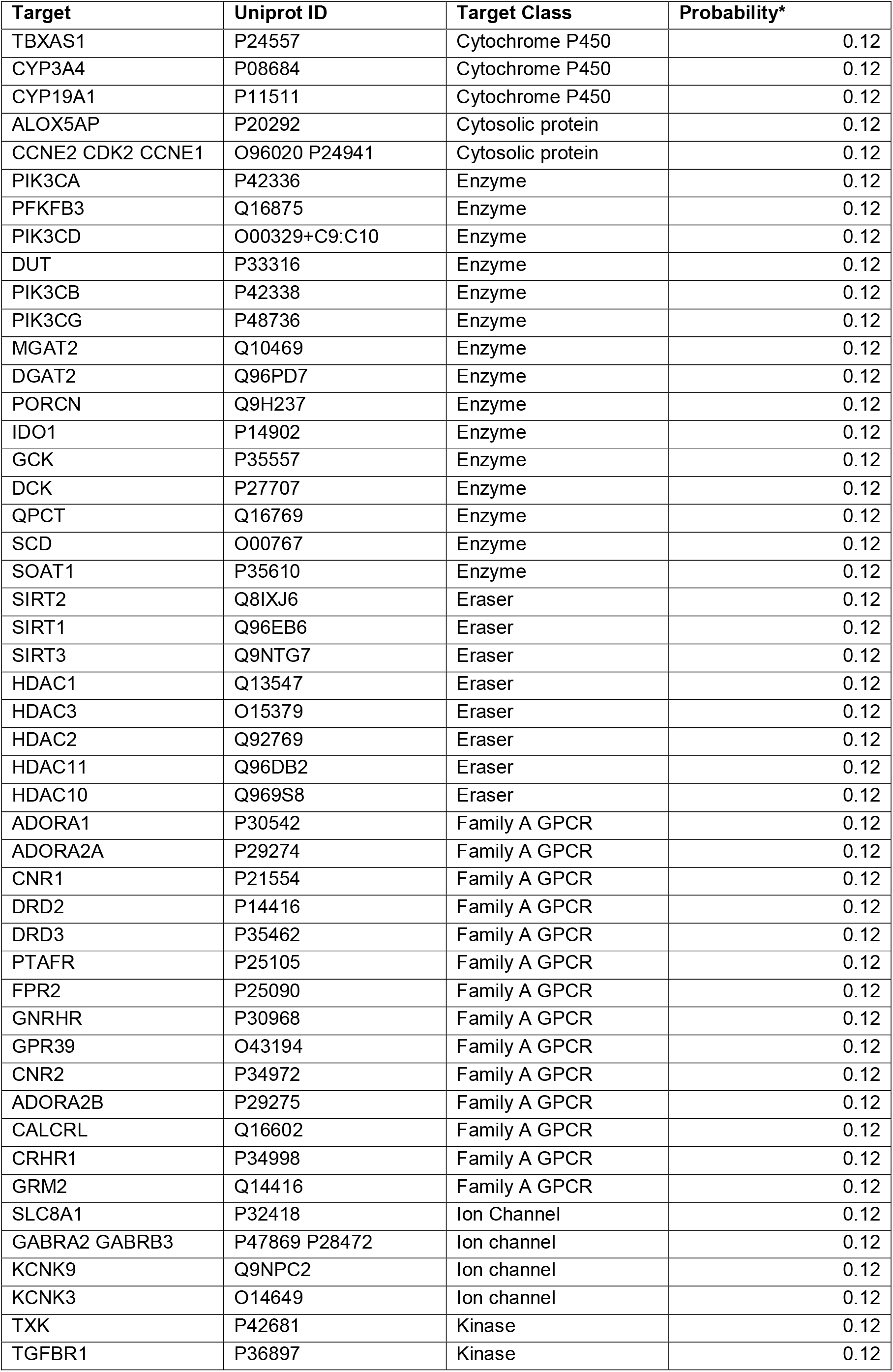
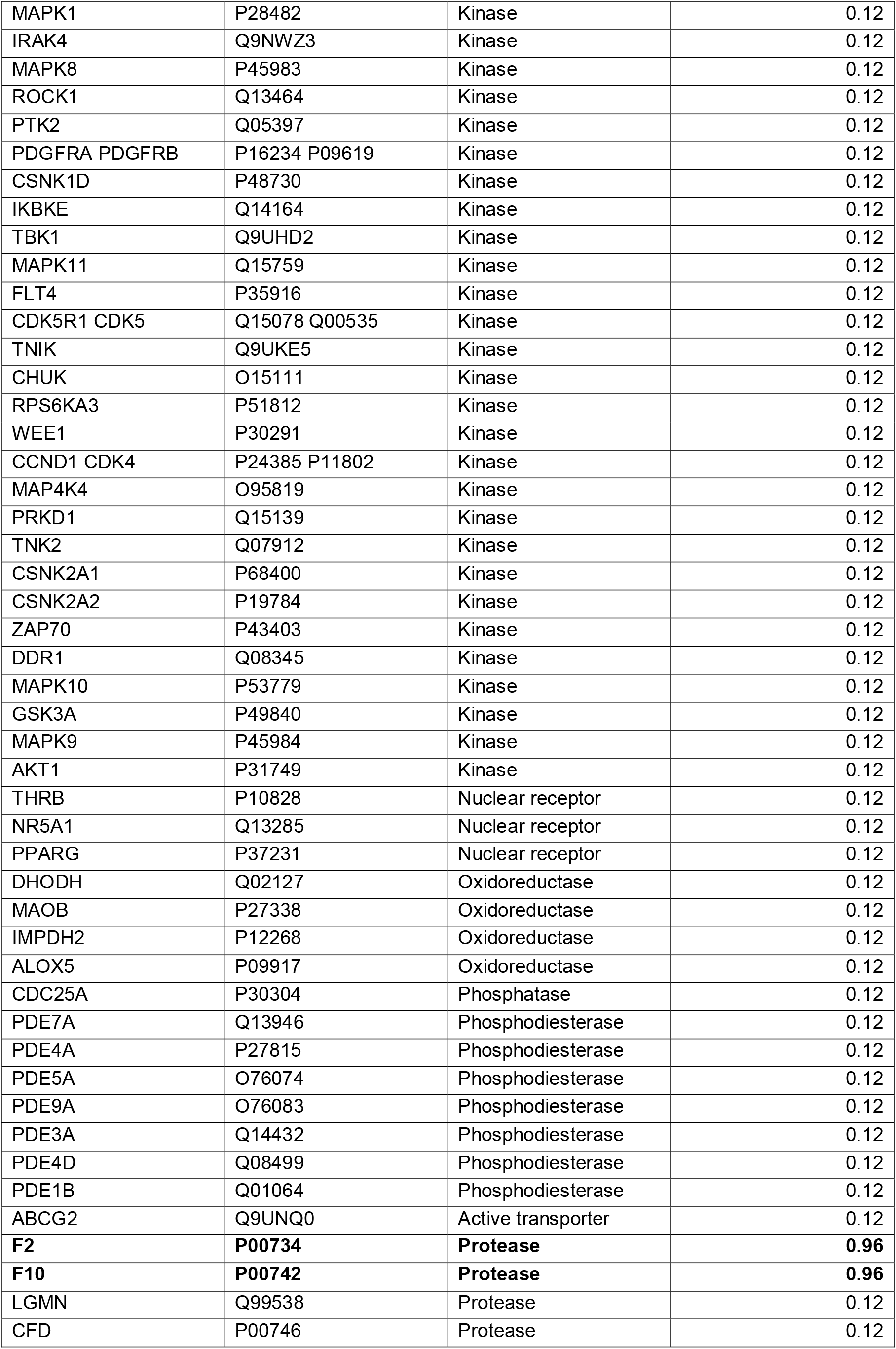
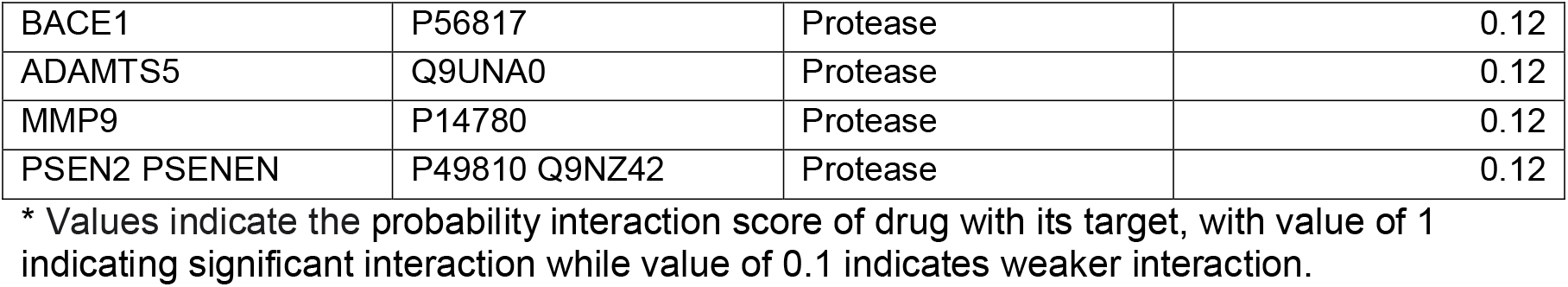
All the potential targets of apixaban in humans.

This study also analysed the target functional categories of each of the anticoagulants, which are summarised in figure 1. The major target categories of warfarin were kinase (29%), enzymes (20%) and family A GPCR (17%) collectively accounting for over 50% of the targets, although the affinity for all these targets was low (probability score < 0.20) (table 2). Similar to warfarin the major target categories of apixaban were also kinase (30%), enzymes (15%) and family A GPCR (14%) (figure 1) all of which showed lower affinity interaction (table 4). About 8% of apixaban targets were proteases among which it had highest affinity (probability score: >0.8) for coagulation factor II and X. The major target categories of dabigatran were proteases (57%) and membrane receptors (13%) (figure 1). Among these targets dabigatran showed high affinity interactions with several proteases and two electrochemical transporters (table 5). Dabigatran was also observed to target quinone reductase 2 with moderate affinity and this is likely to lead to synergistic pharmacological effects with warfarin (which targets quinone reductase 1). Rivaroxaban selectively targeted proteases and was the only DOAC observed to have minimal number of targets (figure 1 and table 3). The major target categories of edoxaban were kinase (29%), protease (21%) and family A GPCR (14%) including low affinity interactions with several off targets including hERG channels (figure 1 and table 6). The major target categories of betrixaban were protease (44%), kinase (17%) and family A GPCR (13%) (figure 1). Betrixaban showed high affinity interaction with coagulation factor X and hERG channels (table 7).

**Table 5:** All the potential targets of dabigatran in humans.

| Target | Uniprot ID | Target Class | Probability* |
| --- | --- | --- | --- |
| **SLC22A2** | **O15244** | **Electrochemical transporter** | **0.99** |
| **SLC47A1** | **Q96FL8** | **Electrochemical transporter** | **0.99** |
| **PLG** | **P00747** | **Protease** | **0.99** |
| **F2** | **P00734** | **Protease** | **0.99** |
| **HPN** | **P05981** | **Protease** | **0.99** |
| **F10** | **P00742** | **Protease** | **0.99** |
| **NQO2** | **P16083** | **Enzyme** | **0.33** |
| TMPRSS15 | P98073 | Protease | 0.11 |
| ITGAV ITGB1 | P06756 | Membrane receptor | 0.11 |
| F3 F7 | P13726 | Protease | 0.11 |
| RPS6KA3 | P51812 | Kinase | 0.11 |
| C3AR1 | Q16581 | Family A GPCR | 0.11 |
| CA12 | O43570 | Lyase | 0.11 |
| ITGA2B | P08514 | Membrane | 0.11 |
| PRSS3 | P35030 | Protease | 0.11 |
| C1S | P09871 | Protease | 0.11 |
| TMPRSS11D | O60235 | Protease | 0.11 |
| ITGA4 | P13612 | Membrane receptor | 0.11 |
| F3 | P13726 | Surface antigen | 0.11 |
| PROC | P04070 | Protease | 0.11 |
| MMP12 | P39900 | Protease | 0.11 |
| MMP8 | P22894 | Protease | 0.11 |
| MMP2 | P08253 | Protease | 0.11 |

\* Values indicate the probability interaction score of drug with its target, with value of 1 indicating significant interaction while value of 0.1 indicates weaker interaction.

**Table 6:**
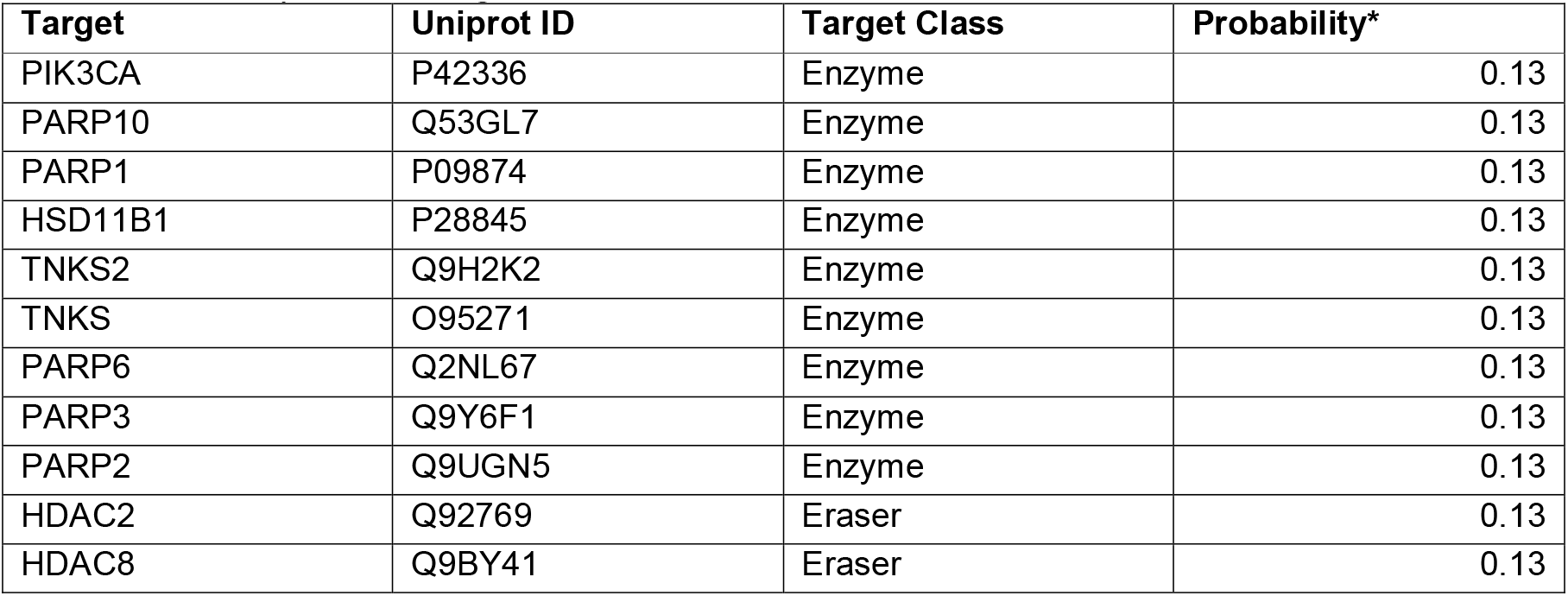
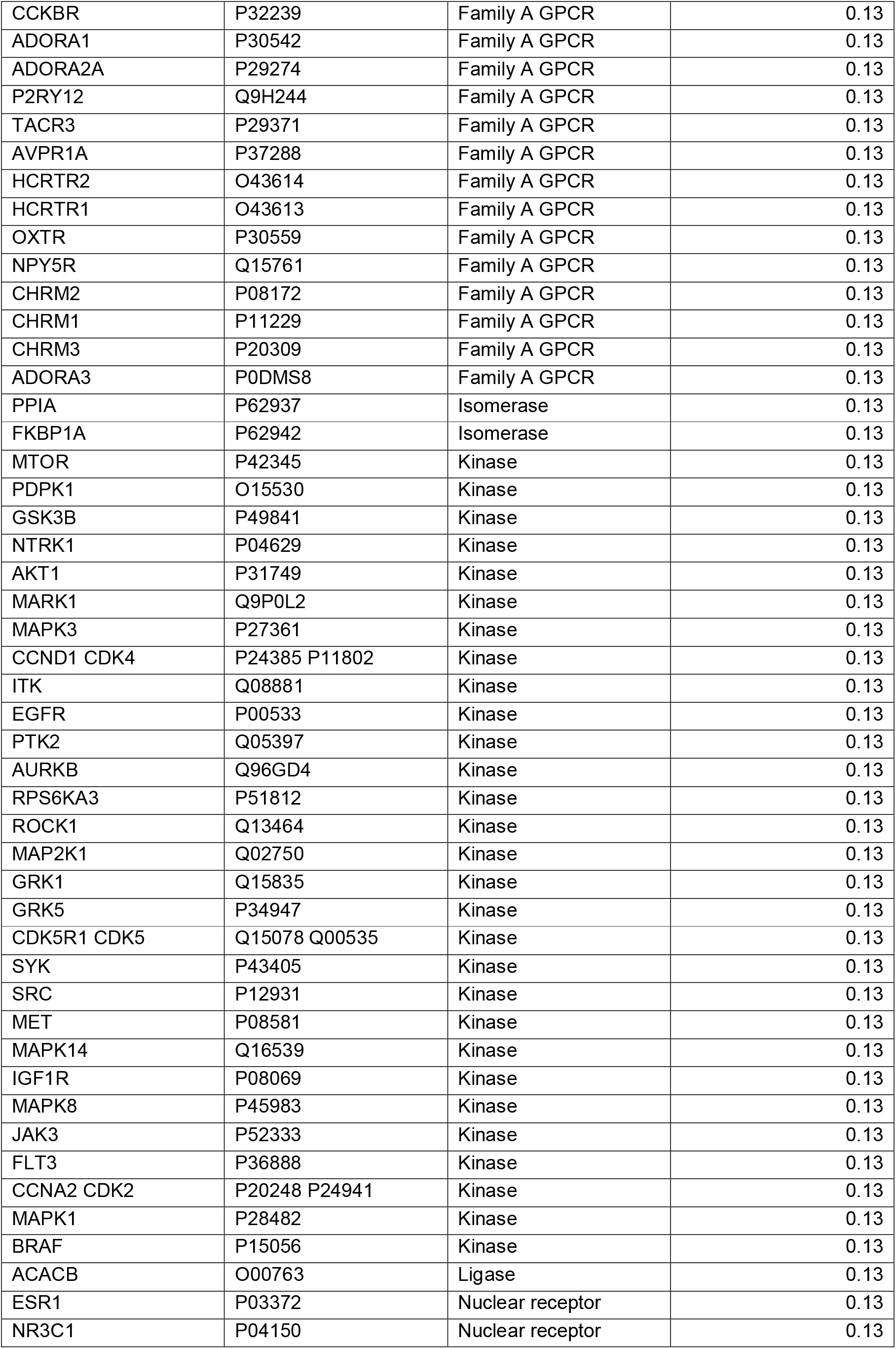
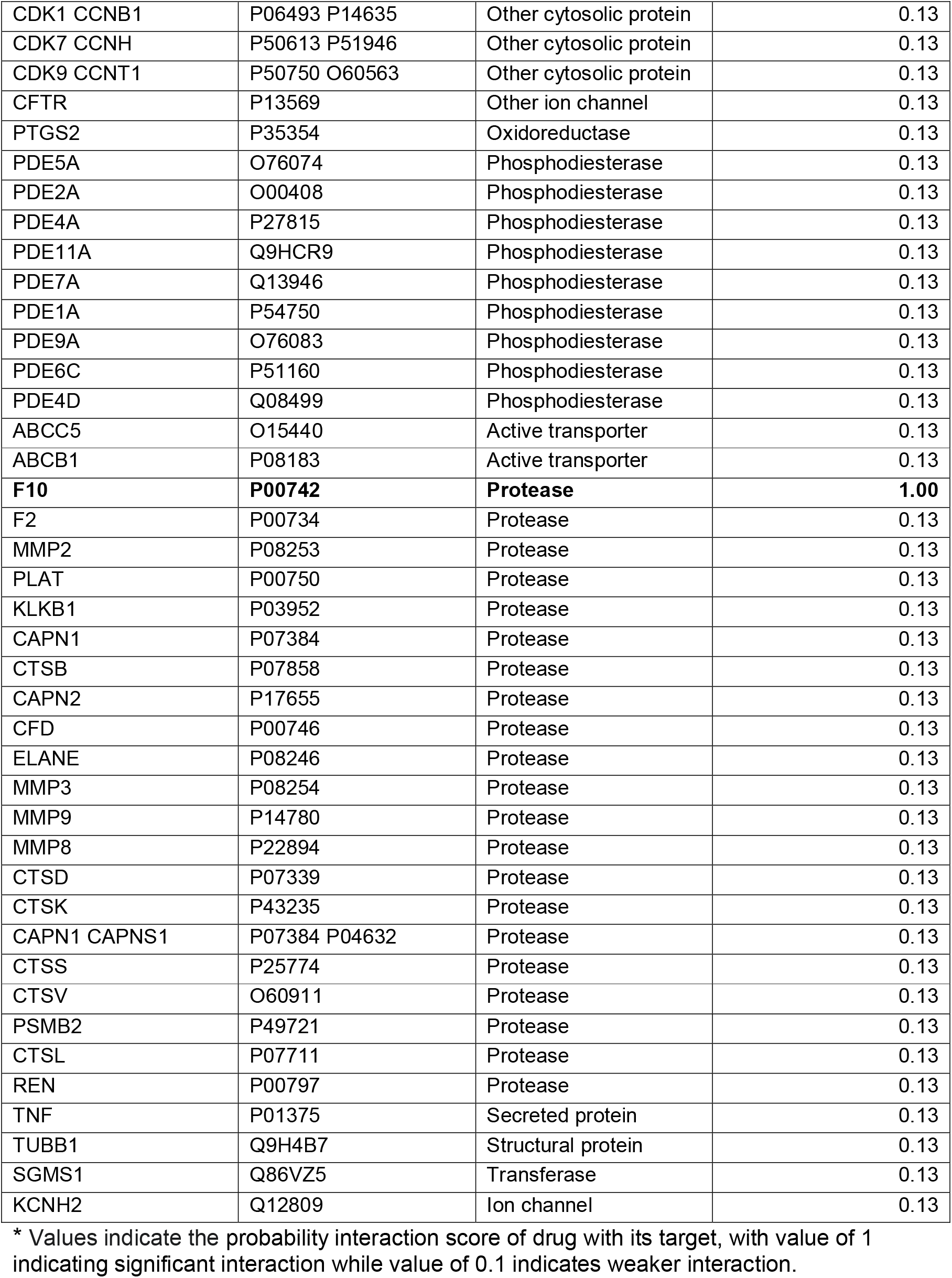
All the potential targets of edoxaban in humans

**Table 7:**
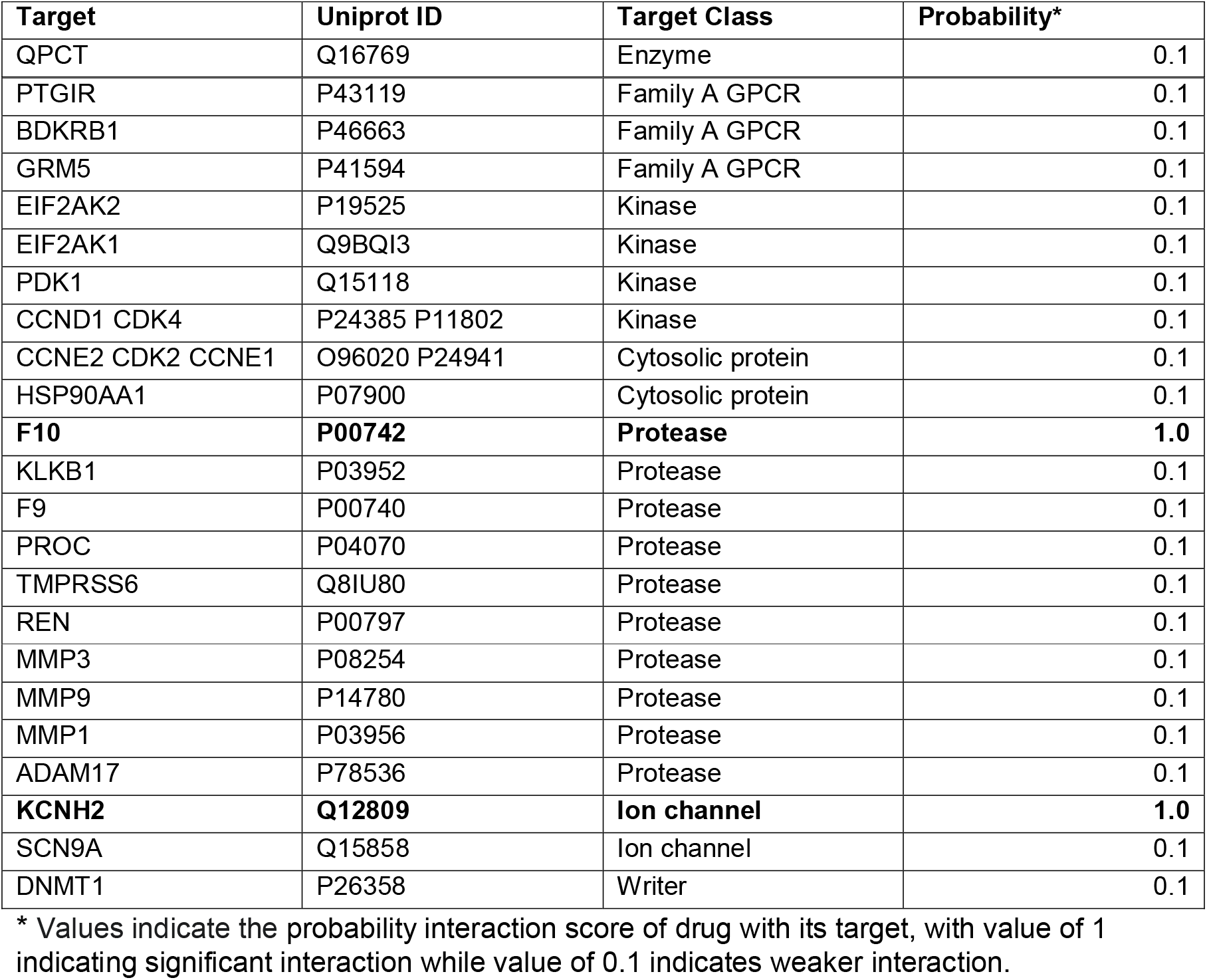
All the potential targets of betrixaban in humans.

To assess the likeliness of off target effect, in this study CA ratio was developed. CA ratio estimates the threshold by which the plasma concentration of the drug is higher than the affinity of the drug to its primary target. Hence drugs with higher CA ratio are highly likely to interact with off targets and produce undesired pharmacodynamics. The CA ratio of warfarin was significantly higher than any of the DOAC (figure 2). Among the DOAC, apixaban had the highest CA ratio especially at higher drug concentration, while apixaban, rivaroxaban, dabigatran and edoxaban showed similar CA ratio for low and medium dose range (figure 2). Betrixaban showed the least CA ratio among the DOAC (figure 2), indicting it being least likely to interact with off target effects. However the high affinity of betrixaban to hERG channels is of concern as highlighted above.

**Figure 2:**
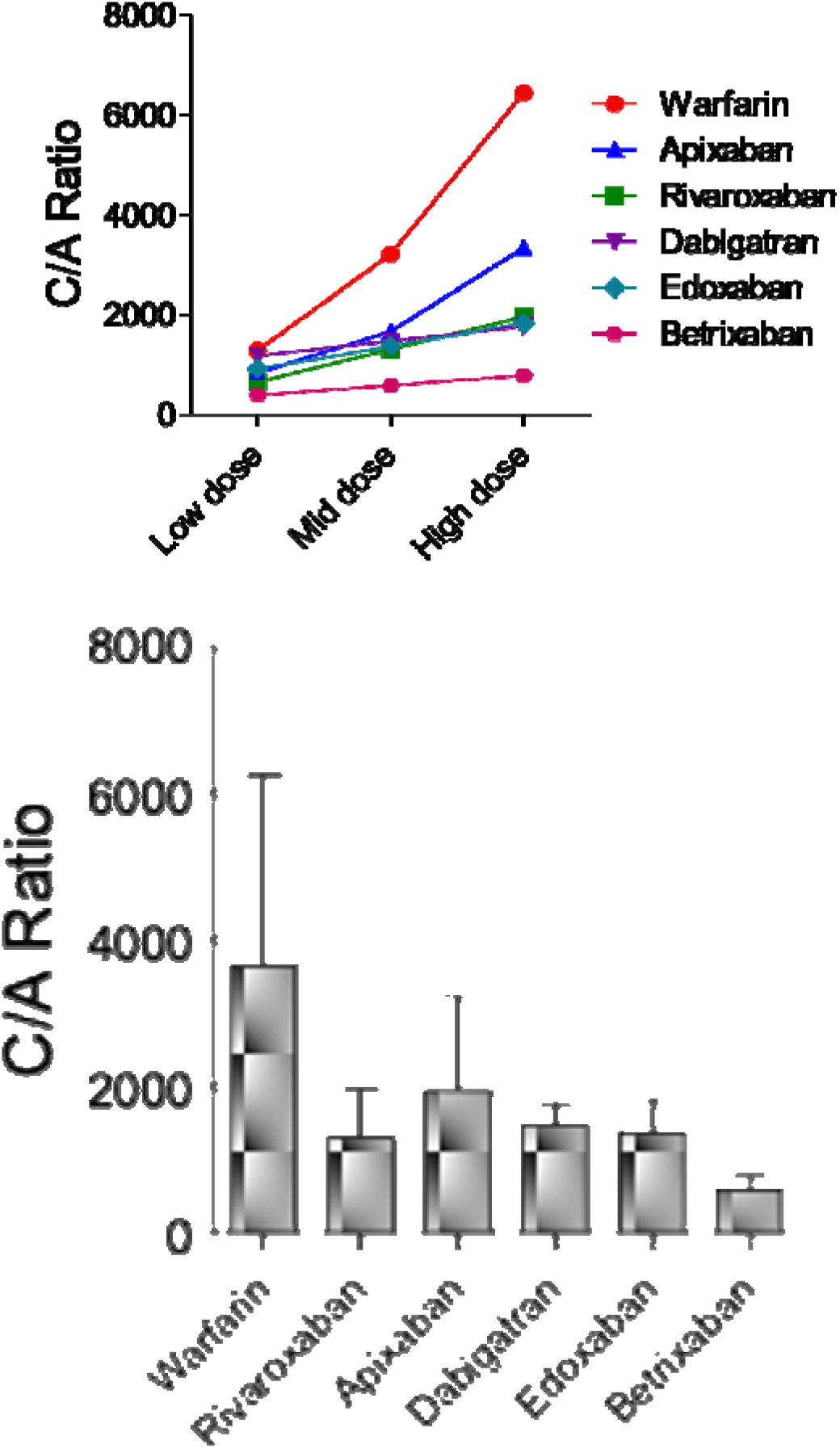
Dose dependent concentration affinity (C/A) ratio of warfarin, apixaban, rivaroxaban, dabigatran, edoxaban, and betrixaban in humans. The bar graphs represents the average C/A ratio. Data is represented as mean ± SD of the C/A ratio values estimated from low, mid and high dose of each of the drugs.

## Discussion

In this analysis apixaban and rivaroxaban were observed to have optimal pharmacological profile for achieving anticoagulation effect. It is also reasonable to conclude that apixaban will have a superior pharmacodynamic outcome compared to rivaroxaban despite its specificity and lower number of off target effects due to the higher affinity of apixaban against factor X and thrombin. These observation are consistent with reports from clinical efficacy trials.^[11, 12, 14]^ Moreover rivaroxaban was observed to off target matriptase with high affinity, which is likely to induce iron deficiency anaemia in patients on chronic use^[19, 20]^ and this is likely to further increase the risk of bleeding. Although apixaban had several off targets, all of these were low affinity interactions, which is least likely to trigger undesired pharmacodynamic effects as long the dosage is adhered to and adequate chrono-pharmacological measures^[21]^ are taken to avoid drug accumulation. The specific higher affinity interactions of apixaban with factor X and thrombin will account for its superior anticoagulation pharmacology. Further the CA ratio of apixaban at low to medium dose range (2 to 5 mg/day) was similar to other DOAC, which together with its low affinity for off targets makes it the DOAC of choice for clinical use over a longer term (greater than 3 months). Considering the selectivity of rivaroxaban to proteases, significantly smaller number of off targets and a lower CA ratio across winder dose range (10 to 30 mg/day), the clinical use of rivaroxaban can be proffered for short term (less than 4 wks). However any long term use of rivaroxaban will require continuous monitoring of haemoglobin (Hb) levels and anaemic status of the patient.

Unlike apixaban or rivaroxaban, dabigatran showed higher affinity to several off targets including quinone reductase 2. Hence dabigatran is likely to show significant undesired pharmacodynamic outcome, including potentiation of warfarin effects.^[22]^ Hence based on the observations from this study the combined use of dabigatran and warfarin should be avoided. The CA ratio of dabigatran was comparable to apixaban and rivaroxaban, hence these DOAC should be proffered over the use of dabigatran. This conclusion is consistent with studies directly comparing these DOAC under clinical setting.^[5, 10-14]^ While edoxaban and betrixaban showed higher affinity to factor X compared to apixaban, both these drugs had several undesirable off targets including the hERG channels. Considering the significantly higher risk of ventricular tachyarrhythmia due to QT prolongation induced by hERG channels, in the opinion of this study use of edoxaban and betrixaban for anticoagulation effects should be avoided.

Several studies have previously reported the superiority of DOAC compared to VKA (warfarin)^[12, 14, 23, 24]^ and this was further evident from the observations in this study. Warfarin showed low affinity interaction with several off targets including its major target (quinone reductase 1) responsible to achieve anticoagulation effects. Further warfarin showed significantly higher CA ratio compared to the DOAC, which increase the likeliness of the warfarin’s off target pharmacodynamics. This observation further supports the wider literature expressing concerns on the relative safety margin of warfarin use.^[25–27]^ Clinically warfarin is also used in combination with DOAC and based on this study it is recommended that only apixaban or rivaroxaban is considered to be used with warfarin when necessary. As mentioned above the use of warfarin with dabigatran should be avoided as this is likely to be associated with higher risk of bleeding and several undesired pharmacodynamic effects.

In summary this study shows the comparative pharmacology of DOAC and VKA and suggests preferential use of apixaban or rivaroxaban due to their superior pharmacodynamic effects and wider safety margin.

## Acknowledgement

Research support from University College Dublin-Seed funding/Output Based Research Support Scheme (R19862, 2019), Royal Society-UK (IES\R2\181067, 2018) and Stemcology (STGY2708, 2020) is acknowledged.

